# Effects of fluorescent glutamate indicators on neurotransmitter diffusion and uptake

**DOI:** 10.1101/2019.12.13.875724

**Authors:** Moritz Armbruster, Chris G. Dulla, Jeffrey S. Diamond

## Abstract

Genetically encoded fluorescent glutamate indicators (iGluSnFRs) enable neurotransmitter release and diffusion to be visualized in intact tissue. Synaptic iGluSnFR signal time courses vary widely depending on experimental conditions and often last 10-100 times longer than the extracellular lifetime of synaptically released glutamate estimated with uptake measurements. iGluSnFR signals typically also decay much more slowly than the unbinding kinetics of the indicator. To resolve these discrepancies, here we have modeled synaptic glutamate diffusion, uptake and iGluSnFR activation to identify factors influencing iGluSnFR signal waveforms. Simulations suggested that iGluSnFR competes with transporters to bind synaptically released glutamate, delaying glutamate uptake. Accordingly, synaptic transporter currents recorded in iGluSnFR-expressing cortical astrocytes were slower than those in control astrocytes. Simulations also suggested that iGluSnFR reduces free glutamate levels in extrasynaptic spaces, likely limiting extrasynaptic receptor activation. iGluSnFR and lower-affinity variants nonetheless provide linear indications of vesicle release, underscoring their value for optical quantal analysis.

## Introduction

Periplasmic binding proteins (PBPs) have been modified to develop genetically encoded biosensors to detect different molecules, including glutamate (de Lorimier et al., 2002). PBPs comprise two domains linked by flexible “hinge” where ligand binding brings the two domains closer together (Quiocho et al., 1997). PBP glutamate indicators are based on GltI, part of *E.Coli.*’s ABC glutamate/aspartate transporter complex; early versions (FLIPE and GluSnFR; Okumoto et al., 2005; Hires et al., 2008) signaled ligand binding via Förster Resonance Energy Transfer (FRET; Fehr et al., 2002) between fluorescent proteins tethered to each PBP domain. Limitations due to low FRET efficiency subsequently were overcome with iGluSnFR, a single-fluorophore sensor with circularly permutated GFP inserted near GltI’s hinge region so that glutamate binding increases GFP fluorescence (Marvin et al., 2013). iGluSnFR variants exhibiting faster glutamate dissociation rates recently have been developed to image glutamate with higher temporal resolution (Helassa et al., 2018; Marvin et al., 2018).

iGluSnFR has been used to detect relative amounts of glutamate release evoked by different physiological stimuli (Borghuis et al., 2013; Yonehara et al., 2013; Armbruster et al., 2016; Franke et al., 2017; Pinky et al., 2018) and to compare the diffusion lifetime of synaptically released glutamate in different brain regions (Pinky et al., 2018). Analogous information has been obtained using synaptically-evoked excitatory amino acid transporter (EAAT)-mediated currents (STCs) recorded in astrocytes (Bergles and Jahr, 1997; Diamond et al., 1998; Luscher et al., 1998; Diamond and Jahr, 2000; Diamond, 2005; Hanson et al., 2015). Although these two approaches can lead to similar conclusions (e.g., Hanson et al., 2015; Armbruster et al., 2016; Pinky et al., 2018), they differ significantly in their response time courses: Synaptically-evoked iGluSnFR-mediated fluorescence signals decay with exponential time courses ranging from ∼20 ms (Marvin et al., 2013; Armbruster et al., 2016) to 100 ms or more (Parsons et al., 2016; Pinky et al., 2018). STCs decay nearly 10 times more rapidly (Bergles and Jahr, 1997; Diamond and Jahr, 2000; Diamond, 2005) and suggest that glutamate is removed from the extracellular space by EAATs in just a few ms (Diamond, 2005). This difference between iGluSnFR response and STC waveforms is evident even in studies employing both techniques under apparently similar experimental conditions (Armbruster et al., 2016).

For iGluSnFR signals and STCs to provide quantitative insights into the dynamics of glutamatergic transmission, the kinetic discrepancies between the two signals must be understood. The STC time course reflects a combination of release asynchrony, transporter kinetics, glutamate diffusion and electrotonic distortion by astrocytic membranes (Bergles and Jahr, 1997; Diamond, 2005). By contrast, the factors determining iGluSnFR response waveforms have not been identified explicitly. Slower iGluSnFR responses do not simply reflect the kinetics of the indicator, as the iGluSnFR dissociation time course is 2-10-fold faster than most response decays. Moreover, iGluSnFR signals are slowed by partial blockade of glutamate transporters (Armbruster et al., 2016; Parsons et al., 2016; Pinky et al., 2018), indicating that they report changes in uptake capacity and glutamate clearance. Whereas STCs reflect the naturally occurring process of glutamate uptake by endogenous transporters, iGluSnFR expression introduces exogenous binding sites into the extracellular milieu. The extent to which glutamate buffering by iGluSnFR may influence glutamate diffusion is not intuitively obvious.

Here, Monte Carlo simulations of glutamate diffusion, uptake and iGluSnFR signaling were performed to explore the mechanisms underlying iGluSnFR signal dynamics. These simulations show that iGluSnFR response time course depends strongly on iGluSnFR expression level. Simulated iGluSnFR responses mimic those reported in the experimental literature only when iGluSnFRs compete with EAATs for glutamate to the extent that iGluSnFR delays glutamate uptake. These predictions were confirmed with electrophysiological recordings from iGluSnFR-expressing astrocytes in cortical slices: STCs recorded in iGluSnFR-expressing astrocytes rose and decayed more slowly that those recorded in control astrocytes expressing tdTomato, indicating that iGluSnFR expression slowed the glutamate uptake time course. We conclude that, although iGluSnFR and STCs provide powerful, complementary indications of glutamate release and clearance, care is required in their interpretation. Our simulations suggest that an ideal glutamate indicator would exhibit a large dynamic range (i.e., ΔF/F_0_) and low expression levels to deliver detectable signals with minimal disruption of glutamate uptake.

## Results

The stochastic behavior of simulated glutamate transporter and iGluSnFR molecules was governed by experimentally derived kinetic models (Figure 1, Supplementary Figure 1). Equilibrium kinetics of simulated EAAT2 (Bergles et al., 2002), iGluSnFR and two iGlu variants (iGlu_f_ and iGlu_u_; Helassa et al., 2018) were examined first by constructing Markov models of each and challenging them with 20-ms applications of glutamate at different concentrations (e.g., Figure 1A). This approach yielded equilibrium dose-response curves (Figure 1B) and affinities (K_D_; Figure 1C, *left*) that matched closely those reported previously (Bergles et al., 2002; Helassa et al., 2018). iGlu molecules exhibited similar activation kinetics but a range of deactivation (unbinding) kinetics that varied inversely with affinity (Figure 1C,D). As the brightness of resting and activated indicator (F_off_ and F_on_, respectively) has been measured (Helassa et al., 2018; Supplementary Figure 1B), iGlu responses also could be expressed in terms of ΔF/F_0_ (i.e., (F_on_-F_off_)/F_off_; Figure 1E).

**Figure 1.**
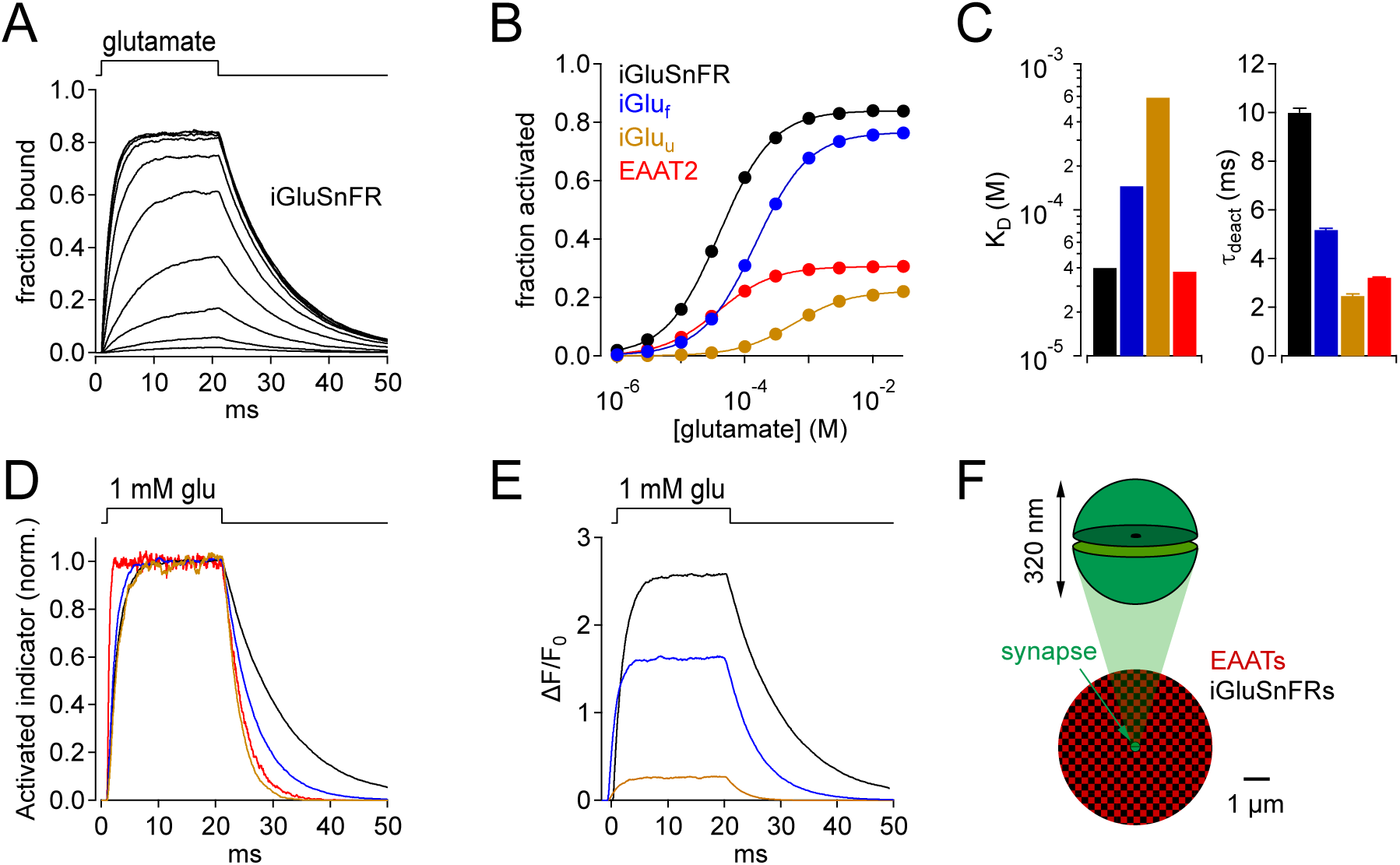
Kinetic properties of glutamate indicators/transporters and simulated synaptic responses. *A*, Simulated iGluSnFR activation by 20-ms applications of glutamate (concentration steps varied logarithmically from 1 μM to 30 mM). *B*, Comparison of simulated glutamate dose-response curves for iGluSnFR (black), iGlu_f_ (blue), iGlu_u_ (gold) and EAAT2 (red). Color scheme applies to the entire figure. *C*, Simulated equilibrium affinities (K_D_, *left*) and deactivation time constants (*right*) for iGlus and EAAT2. *D*, Activation of iGlus and EAAT2 by 1 mM glutamate, normalized and superimposed to compare activation and deactivation kinetics. *E*, Responses of iGlus to 1 mM glutamate, scaled according to their background fluorescence and change in fluorescence upon activation (i.e., ΔF/F_0_ ≍ (F_on_-F_off_)/F_off_; see Supplementary Figure 1; Helassa et al., 2018). *F*, schematic diagrams of simulated synaptic structure (*top*) and surrounding extracellular space (*bottom*).

### Stochastic model of glutamate diffusion, uptake and iGluSnFR activation

Monte Carlo simulations of glutamate release from a single synapse (Diamond, 2005; see Methods) comprised pre- and postsynaptic hemispherical compartments (320-nm diameter; Schikorski and Stevens, 1997; Ventura and Harris, 1999) separated by a 20-nm cleft and surrounded by a three-dimensional, isotropic, abstract representation of extracellular space (Rusakov and Kullmann, 1998) populated with EAAT and iGluSnFR molecules at specified concentrations (Figure 1F). EAATs are expressed in astrocytic membranes at high densities (> 10^4^ molecules per μm^2^) that, adjusted for membrane density and extracellular volume fraction, correspond to effective concentrations of 140-330 μM in the extracellular space of hippocampal and cerebellar neuropil (Lehre and Danbolt, 1998). Accordingly, the time course of glutamate uptake in adult CA1 hippocampal astrocytes is well modeled with an active EAAT concentration of about 100 μM (Diamond, 2005), the value used here. iGluSnFR concentrations have not been measured but, because they may vary widely depending on factors influencing expression, we simulated a large range (1-3000 μM).

5000 glutamate molecules were released at the center of the synaptic cleft, with each individual glutamate molecule undertaking a random walk slowed by the tortuosity of the extracellular space (Nicholson and Sykova, 1998; Rusakov and Kullmann, 1998; Nielsen et al., 2004; Diamond, 2005). The extrasynaptic space was populated evenly with completely overlapping distributions of EAATs and iGluSnFRs. Because glial and neuronal membranes are so closely apposed in synaptic neuropil (Mishchenko et al., 2010), the large majority of glial EAATs likely must compete with iGluSnFRs for glutamate, regardless of whether iGluSnFR is expressed pan-neuronally or in glia (Armbruster et al., 2016). Additional experiments, presented below (Figure 8), examined other extrasynaptic expression patterns of EAATs and iGluSnFRs.

### Competition between iGluSnFRs and EAATs slows uptake and iGluSnFR activation time courses

When EAATs (100 μM) and iGluSnFRs (300 μM) were co-localized in extrasynaptic space, simulated glutamate release activated iGluSnFR with a time course that reached a peak in about 5 ms and decayed with an exponential time course (τ = 27 ms; Figure 2A). This decay was significantly slower than iGluSnFR’s deactivation time constant (τ = 10 ms; Figure 1C, *right*), suggesting that iGluSnFR interacted with extrasynaptic glutamate over a prolonged period. Consistent with this, the time course of glutamate uptake from the extracellular space was slowed in the presence of iGluSnFR (Figure 2B). This slowing was also evident in the simulated STC (Figure 2B, inset), which reflects electrogenic state transitions within EAATs upon binding and transporting glutamate (Bergles et al., 2002; see Methods). The time courses of both the iGluSnFR response and glutamate uptake were prolonged by higher iGluSnFR concentrations (Figure 2C,D), suggesting that iGluSnFR buffers glutamate diffusion and delays its uptake. At very low iGluSnFR levels (e.g., 1 μM), the uptake time course closely approximated the room temperature control in the absence of iGluSnFR (τ ∼ 4 ms; Diamond, 2005), and iGluSnFR activation decayed at a rate approaching iGluSnFR deactivation (Figure 2D). At higher iGluSnFR concentrations, however, both time constants increased and converged; the highest iGluSnFR levels tested gave rise to time constants that exceeded 100 ms (Figure 2D), similar to slower published iGluSnFR signal time courses (Parsons et al., 2016; Pinky et al., 2018). Although iGluSnFR extended the extracellular lifetime of glutamate, it did not affect the distance that glutamate diffused prior to being taken up by transporters (Figure 2E,F), consistent with iGluSnFR’s role as a stationary buffer. At higher expression levels, each glutamate molecule bound to iGluSnFR multiple times before it was able to bind a transporter (Figure 2F). (Each glutamate molecule typically bound a transporter only once, as simulated EAATs transported glutamate with 90% efficiency.) These results indicate that strong iGluSnFR buffering slows glutamate uptake.

**Figure 2.**
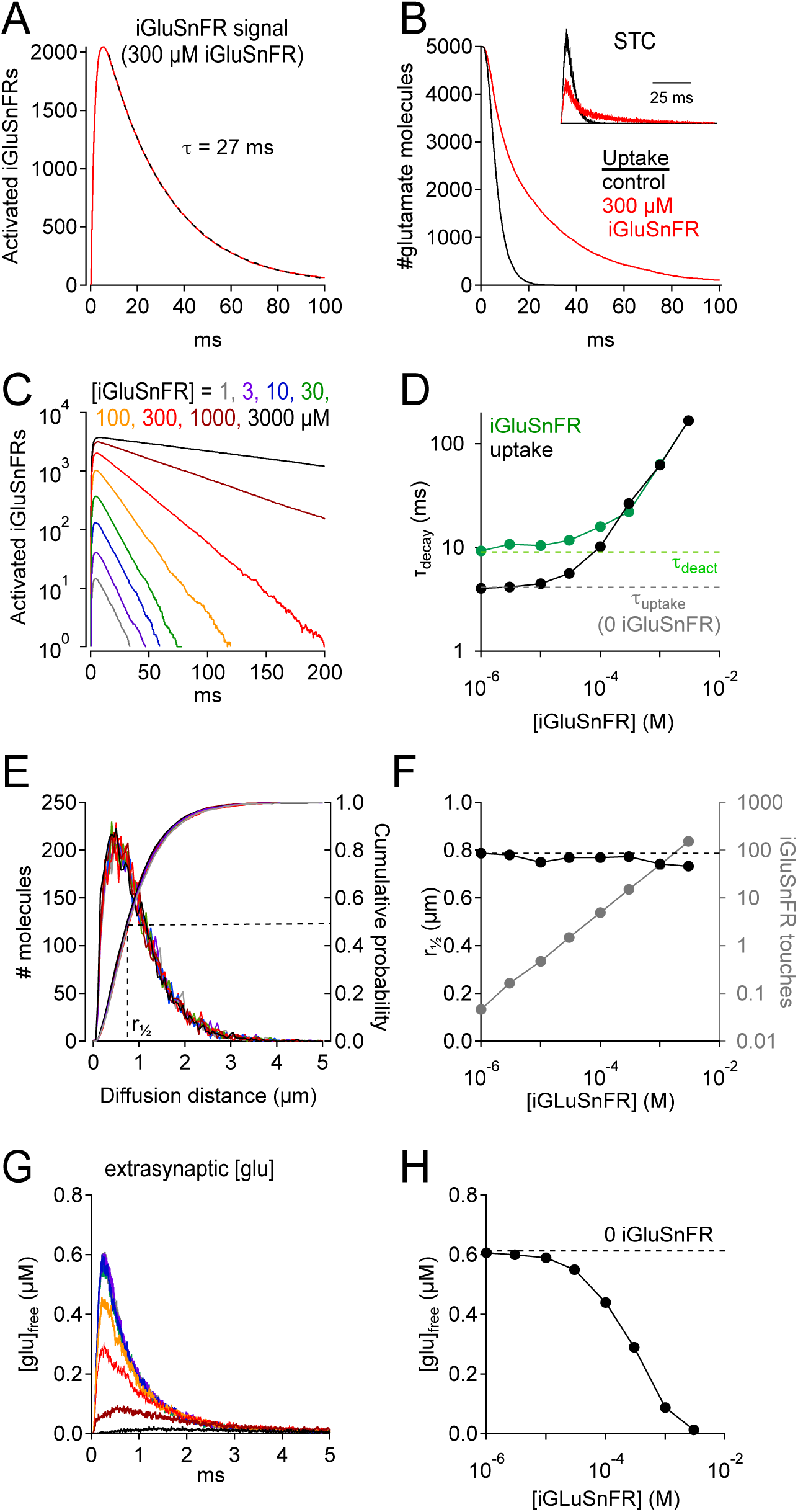
iGluSnFR can distort fluorescence and glutamate clearance time courses. *A*, Simulated iGluSnFR activation elicited by the release of 5000 glutamate molecules from the center of the cleft in the scheme depicted in Figure 1*F*. Dashed line indicates exponential fit to the response decay. Activation measured over a region of interest (ROI, radius = 10 μm) centered about the synapse. *B*, Simulated time course of glutamate uptake in the absence of iGluSnFR (black) and in the presence of 300 μM iGluSnFR (red). *Inset*, simulated STCs in the presence (red) and absence (black) of 300 μM iGluSnFR. STCs are typically inward (negative) currents but are inverted here for simplicity. *C*, The concentration of iGluSnFR influences its activation time course. A semi-log plot shows that the exponential decay slows as [iGluSnFR] increases (ROI radius = 10 μm). *D*, Summarized data from *C* (green). Glutamate uptake (black) is also slowed by [iGluSnFR]. *E*, iGluSnFR buffering does not affect the distance that glutamate diffuses prior to being taken up by EAATs. Same color scheme as in *C*. Distance measured from the center of the synaptic cleft. *F*, Summarized data from *E*. Gray trace shows the average number of times each glutamate molecule binds to iGluSnFR prior to being taken up by an EAAT. *G*, iGluSnFR buffering reduces the extrasynaptic concentration of free glutamate. Traces show the glutamate concentration in a spherical shell 400-500 nm from the release point. *H*, Summarized data from *G*.

If iGluSnFR buffered synaptically released glutamate, it should reduce the concentration of free neurotransmitter in the extrasynaptic space. To test this, we measured the simulated free (unbound, untransported) glutamate concentration in a concentric spherical shell 400-500 nm from the center of the synapse (Figure 2G), a distance approximating the average distance between CA1 hippocampal synapses (465 nm; Rusakov and Kullmann, 1998). The peak concentration of free glutamate in this shell decreased at higher iGluSnFR levels (Figure 2G,H), suggesting that iGluSnFR may reduce extrasynaptic receptor activation and/or glutamate spillover between synapses.

### iGluSnFR expression slows STCs in cortical astrocytes

The simulations presented so far suggest that iGluSnFR delays glutamate uptake by competing with EAATs and predict that iGluSnFR expression would slow the STC time course in astrocytes. To test this, we recorded STCs in cortical astrocytes from mice expressing either iGluSnFR or tdTomato under control of a glial-specific promoter (GFAP; Figure 3). Consistent with the model’s predictions, STCs in iGluSnFR^+^ astrocytes rose and decayed more slowly than those in tdTomato^+^ astrocytes (t_rise_ (10-90%): 6.6 ± 1.8 ms, n=23 vs. 5.0 ± 2.1 ms, n=15, *t*(26.1)=2.42, *p*=0.023, *t*-test; τ_decay_: 19.2 ± 5.6 ms, n=23 vs. 16.0 ± 3.6 ms, n=15, *t*(36.0)=2.12, *p*=0.041; Figure 3A-D). This was not due to any apparent changes in astrocyte intrinsic electrical properties: iGluSnFR^+^ and tdTomato^+^ astrocytes exhibited similar input resistance (iGluSnFR: 2.9 ± 2.6 MΩ; n=22; tdTomato: 3.6 ± 2.6 MΩ, n=16; *t*(32)=-0.75, *p*=0.46, *t*-test), and iGluSnFR^+^ astrocytes actually rested at slightly more hyperpolarized potentials (−73.3 ± 3.0 mV, n=22 vs. - 70.0 ± 4.0 mV, n=15; *t*(24)=-2.73, *p*=0.011, *t*-test; Figure 3E) that, if anything, would increase slightly the efficacy of glutamate uptake (Wadiche et al., 1995). These changes in STC waveform occurred under experimental conditions that yielded iGluSnFR decay time courses (τ = 21.1 ± 4.9 ms, n=10; Figure 3F) that are near the fast end of the range of published iGluSnFR signals and correspond to responses simulated with 300 μM iGluSnFR (Figure 2D).

**Figure 3.**
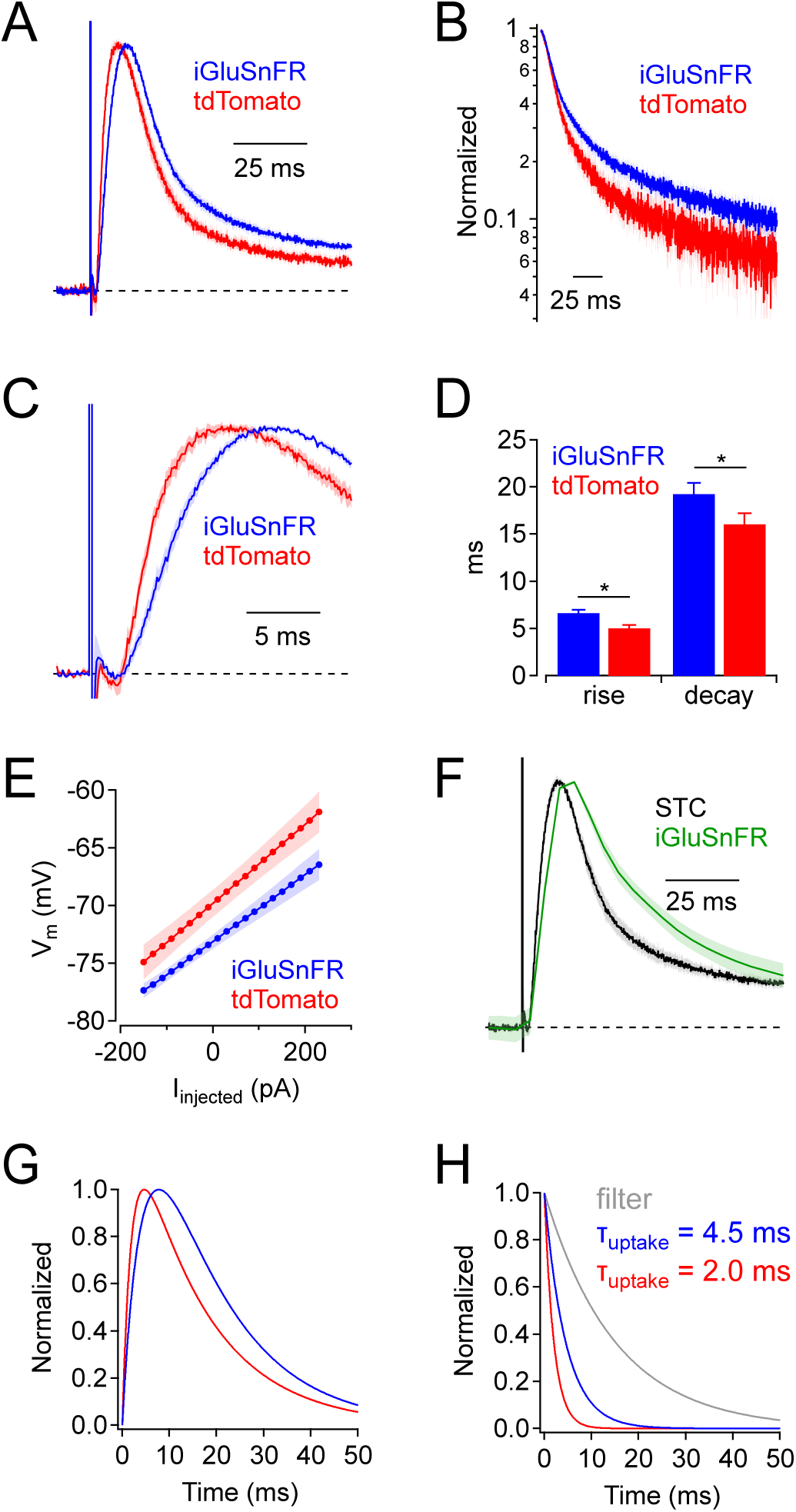
iGluSnFR expression slows uptake by cortical astrocytes. *A*, Synaptic transporter currents (STCs) recorded in cortical astrocytes expressing either iGluSnFR (blue) or tdTomato (red). Traces indicate average responses (mean ± SEM; iGluSnFR: n = 21 cells; tdTomato: n = 15 cells), normalized in amplitude. *B*, Responses in *A*, plotted on a semi-log scale. *C*, Rising phases (± SEM) of the responses in A, plotted on an expanded time scale. *D*, Summary data showing that STC rise and decay were slower in iGluSnFR^+^ astrocytes. * indicates p < 0.05. *E*, Astrocyte V_m_ (mean ± SEM), as a function of injected current, shows that iGluSnFR^+^ astrocytes rested at slightly more hyperpolarized potentials compared to tdTomato^+^ astrocytes (*t*(34.6)=-2.73, *p*=0.011, *t*-test). Input resistances (indicated by the slope of the relation) was not different in the two groups (*t*(32.0)=-0.75, *p*=0.46, *t*-test). *F*, STCs and iGluSnFR signals measured in the same experiments (mean ± SEM, n = 10 cells). *G*, Simulated STC waveforms corresponding to average responses in iGluSnFR^+^ (blue) and tdTomato^+^ (red) astrocytes. *H*, Waveforms used to derive STCs in *G*. In each case a clearance time course (red or blue) was convolved with a filter waveform (gray). This simple example demonstrates how even subtle differences in STC time course can reflect substantial differences in glutamate clearance time course (Diamond, 2005).

At first glance, these differences in STC time course might appear relatively subtle, but they likely indicate substantial modifications of the glutamate clearance time course. Previous analyses of STCs in CA1 hippocampal astrocytes showed that the STC waveform reflects the time course of glutamate uptake filtered primarily by the electronic properties of the astrocyte (Diamond, 2005). This filtering slows both the rise and decay of the STC and obscures the actual time course of glutamate clearance. Consequently, even small changes in STC waveform likely indicate significant changes in glutamate clearance (e.g., Figure 3G,H). Note that simulated STCs and derived uptake times do not reflect any electrotonic distortion or release asynchrony.

### iGluSnFR signal time course and ΔF/F_*0*_ *depends on imaging volume*

iGluSnFR enables glutamate to be imaged over a range of spatial scales, from a <1 μm synapse to an entire brain region (Marvin et al., 2013). Glutamate clearance from a synapse is driven primarily by diffusion down a locally steep concentration gradient (Wahl et al., 1996; Diamond and Jahr, 1997; Barbour, 2001), so that the fractional reduction in glutamate concentration is fastest close to the point of release (Barbour and Hausser, 1997). Accordingly, simulations indicated that iGluSnFR activation signals were faster when measured over a smaller spherical region of interest (ROI) surrounding the release site (Figure 4A,B), consistent with experimental results (e.g., Marvin et al., 2013). This volume-dependent effect was greater at higher iGluSnFR concentrations because stronger buffering prolonged further the extrasynaptic lifetime of glutamate (Figure 4B). At higher expression levels, iGluSnFR bound simultaneously a significant fraction of synaptically released glutamate, approaching levels limited by the kinetic maximum probability of fluorescence (P_max_, analogous to the maximal open probability of an ion channel; Figure 4C and 1B). Consequently, the iGluSnFR activation coefficient of variability (CV = σ/mean) decreased at higher iGluSnFR concentrations (Figure 4D).

**Figure 4.**
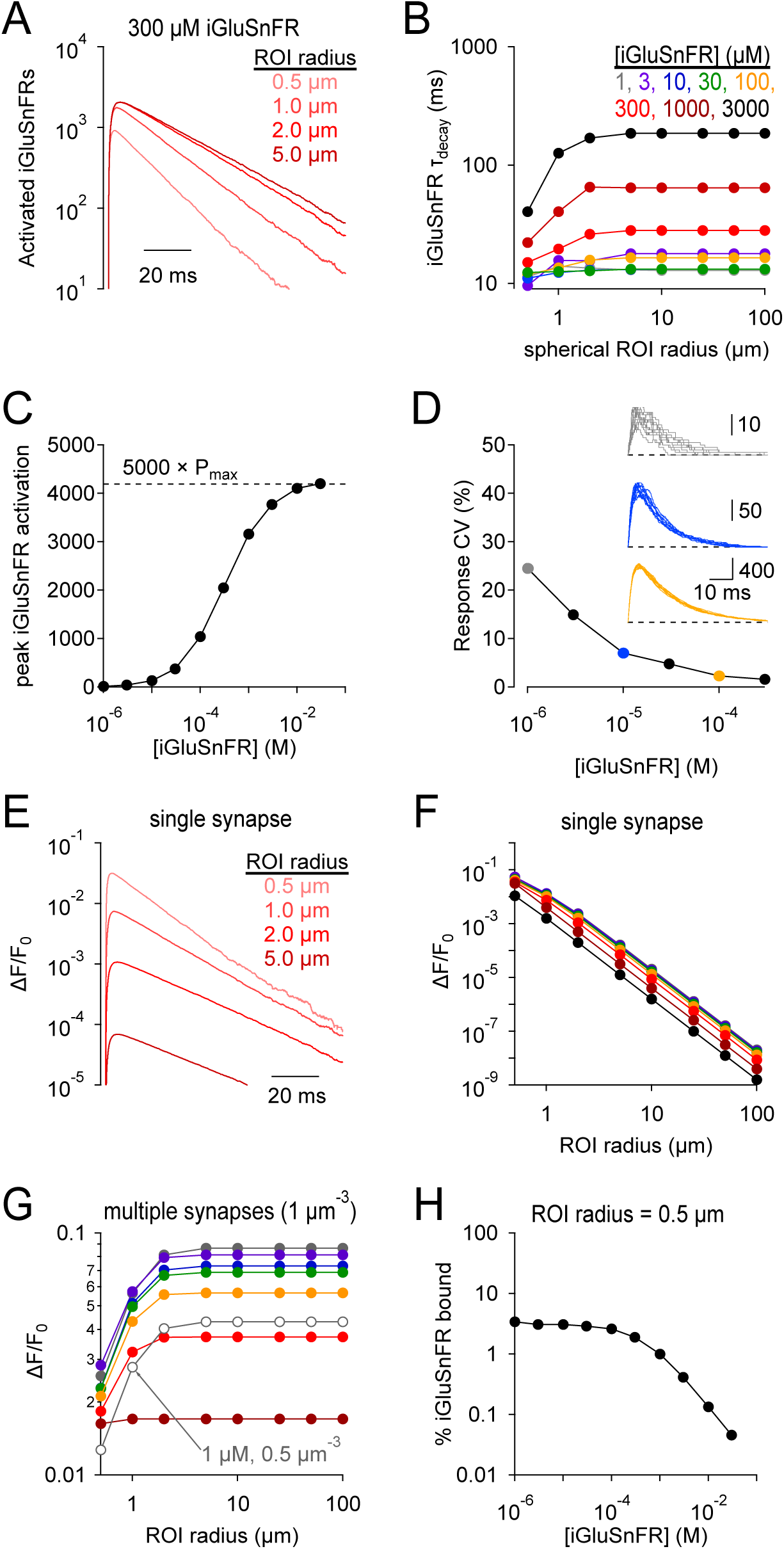
iGluSnFR signal time course and SNR depends on the imaging volume. *A*, iGluSnFR (300 μM) signals measured across different spherical regions of interest (ROIs). *B*, Summary data shows that the dependence on ROI volume is greater at higher iGluSnFR concentrations. *C*, Peak iGluSnFR response to the release of 5000 glutamate molecules as a function of iGluSnFR concentration. Dashed line indicates maximal signal based on maximal occupancy (see Figure 1B). *D*, iGluSnFR signal variability decreases with indicator concentration. *Inset*, individual responses at three different iGluSnFR concentrations (gray, 1 μM; blue, 10 μM; orange, 100 μM). *E-F*, iGluSnFR single-synapse response signal (ΔF/F_0_) depends on indicator concentration (*E*) and ROI dimensions (*F*). *G*, iGluSnFR compound ΔF/F_0_ responses depend on ROI dimensions and the density of activated synapses. *H*, When iGluSnFR was evenly sampled throughout even a small volume, only a small fraction of the indicator was activated by glutamate. Laser line scanning, by contrast, yields higher ΔF/F_0_ values (Helassa et al., 2018).

The brightness of iGluSnFR has been measured in the unbound and activated states (e.g., Helassa et al., 2018), allowing simulated iGluSnFR activation to be expressed in terms of fluorescence, typically reported as the change in fluorescence relative to resting levels (ΔF/F_0_; Figure 4E). Because inactive iGluSnFR in extrasynaptic tissue contributes to F_0_, the single synapse ΔF/F_0_ decreased dramatically over larger imaging volumes (Figure 4E,F), underscoring the necessity of highly localized laser scanning fluorescence microscopy for examining iGluSnFR signals at single synapses (Helassa et al., 2018; Marvin et al., 2018). When compound iGluSnFR responses are recorded from many synapses simultaneously, often using wide-field imaging techniques (e.g., Figure 3), both the number of imaged synapses and the background fluorescence increase proportionally to imaging volume, so that ΔF/F_0_ does not vary greatly over most ROI dimensions (Figure 4G). Note that ΔF/F_0_ values decreased as iGluSnFR expression level increased (Figure 4F,G). This effect appears due primarily to the increasing F_0_, not a significant decrease in the fraction of iGluSnFR bound, because even at very low iGluSnFR concentrations glutamate bound only a small fraction of iGluSnFR within a 0.5 μm-radius sphere surrounding the synapse (Figure 4H). These results point to a somewhat counterintuitive conclusion that increased iGluSnFR expression may, in many experimental conditions, actually decrease ΔF/F_0_ values of synaptic responses.

### Effects of blocking EAATs on iGluSnFR signals

The rate at which glutamate is taken up from the extracellular space depends critically on EAAT expression levels in astroglia (Bergles and Jahr, 1997; Diamond and Jahr, 2000; Diamond, 2005; Thomas et al., 2011). In the hippocampus, the EAAT2 subtype constitutes ∼80% of glial glutamate transporters, and the remaining 20% are EAAT1 (Lehre and Danbolt, 1998). A recent study reported large differences between the effects of an EAAT2-selective antagonist and complete EAAT blockade by a pan-EAAT antagonist on iGluSnFR response time course in hippocampus, leading the authors to suggest that EAAT1 may play a particularly large role in glutamate uptake (Pinky et al., 2018). By contrast, STC recordings suggest that blocking EAAT2 slows glutamate uptake by about five-fold (Diamond and Jahr, 2000; Diamond, 2005), consistent with the relative expression levels of the two transporter subtypes (Lehre and Danbolt, 1998). To examine this discrepancy, we simulated the effects of changing transporter density on iGluSnFR activation time course (Figure 5). The localization and kinetic properties of EAATs remained the same: only transporter density was changed. The apparent effects of simulated EAAT blockade depended dramatically on the imaging volume. Over a 1-μm radius ROI, the effects on iGluSnFR time course of removing 80% or 100% of the transporters was relatively minor (Figure 5A), because the glutamate concentration time course over that small spatial scale is dominated by diffusion, not glutamate uptake (Diamond and Jahr, 1997). By contrast, iGluSnFR responses measured over a 5 μm-radius ROI were slowed dramatically by reducing EAAT density, with a particularly large difference observed between 80% and 100% blockade (Figure 5B). Note that a 5-μm-radius ROI reported accurately the (iGluSnFR-buffered) time course of glutamate clearance across a range of EAAT levels (Figure 5F). These results suggest that a) using iGluSnFR to evaluate manipulations of glutamate diffusion and uptake requires careful consideration of imaging parameters, and b) even a small fraction of expressed EAATs clears glutamate relatively quickly, such that reducing EAATs from control to 20% exerts less dramatic effects on iGluSnFR signals than reducing EAATs from 20% to zero, regardless of EAAT subtype (Figure 5B; Pinky et al., 2018).

**Figure 5.**
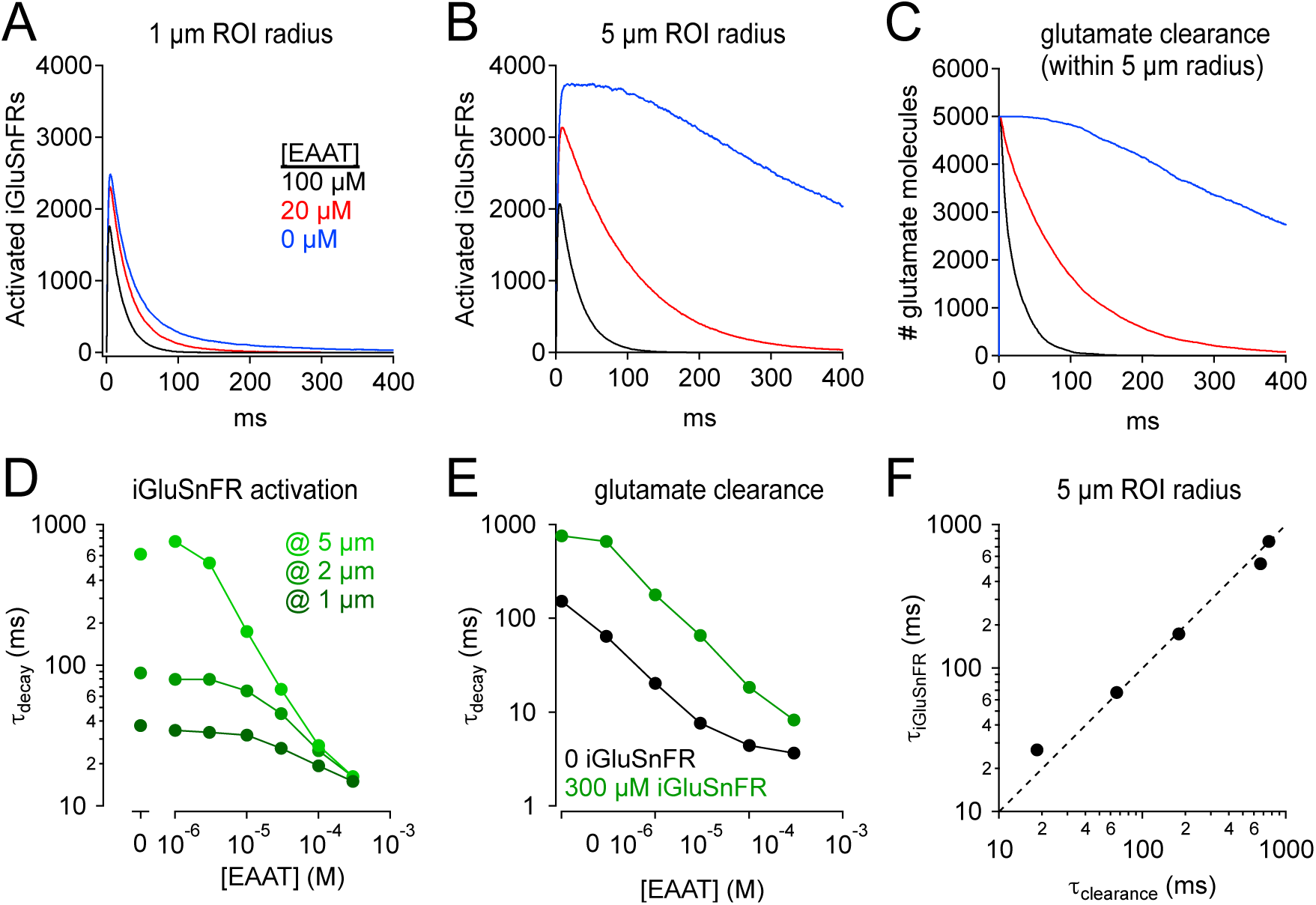
iGluSnFR distorts effects of varying uptake capacity. *A*, iGluSnFR responses (300 μM iGluSnFR) simulated in three different EAAT concentrations. In the immediate vicinity of the synapse (1 μm ROI radius), reducing uptake capacity has little effect on iGluSnFR signal amplitude or time course. *B*, As in *A*, but over a larger imaging volume (5 μm ROI radius), which exaggerates the effects of reducing uptake capacity. *C*, Time course of glutamate clearance (uptake and diffusion beyond a 5 μm radius) in the conditions shown in *A* and *B. D*, Summary graph showing the effects of EAAT concentration and ROI dimensions on the exponential decay time course of iGluSnFR activation. *E*, The time course of glutamate clearance in the absence of indicator (black) and in the presence of 300 μm iGluSnFR (green). F, The iGluSnFR signal measured across a 5-μm-radius ROI accurately reports the (modified) time course of glutamate clearance across a range of EAAT concentrations.

### Lower-affinity iGluSnFR variants do not eliminate buffering artifacts

Recent molecular modifications of iGluSnFR have produced variants that exhibit lower affinity for glutamate, typically by speeding glutamate unbinding (Figure 1B-D, Supplementary Figure 1; Helassa et al., 2018; Marvin et al., 2018). These variants may therefore provide the faster fluorescent signals required to image glutamate dynamics accurately, potentially at individual synapses, and resolve individual responses during high-frequency stimulation (Helassa et al., 2018; Marvin et al., 2018). To test how decreased affinity might influence glutamate signaling, iGluSnFR was replaced in the model by one of two lower-affinity variants, iGlu_u_ or iGlu_f_, with well-characterized kinetic properties (Helassa et al., 2018; Figure 6). Both indicators gave rise to faster responses than did iGluSnFR (cf. Figure 6A,C and Figure 2C) but both produced signal time courses that varied with indicator concentration and imaging volume (Figure 6A-D).

**Figure 6.**
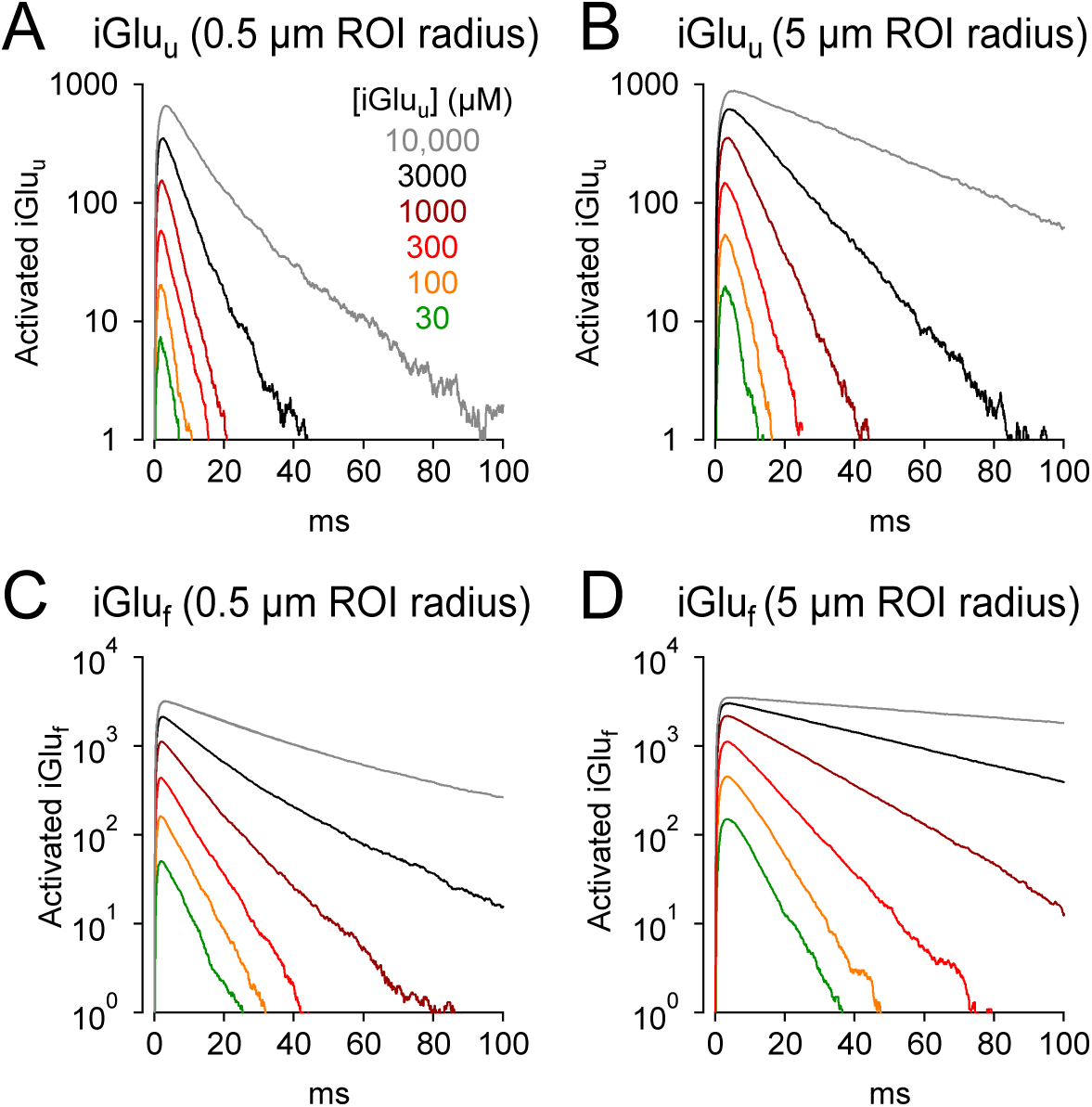
Faster glutamate indicators do not eliminate distortion. *A*, iGlu_u_ signals (0.5 μm ROI radius) simulated with different concentrations of the lower-affinity indicator. *B*, As in *A* but over a 5 μm ROI radius. *C* and *D*, As in *A* and *B* but with iGlu_f_.

### iGluSnFR provides linear indication of glutamate release

iGluSnFR and its variants are potentially valuable tools for comparing relative amounts of glutamate released under different experimental conditions. STCs provide accurately proportionate indications of glutamate release (Diamond et al., 1998; Luscher et al., 1998), but similar calibrations of iGluSnFRs have not been performed. To test this, we simulated coincident neurotransmitter release from variable numbers of synapses arranged at different densities (Figure 7). The diffusion medium included 100 μM EAAT and 300 μM iGluSnFR, a combination that produced consistent, sizeable iGluSnFR signals at individual synapses (Figure 2A) and approximated experimentally observed iGluSnFR time courses (Figure 3F). In these multi-synapse simulations, the 30×30×30-μm^3^ diffusion space was mapped in Cartesian coordinates and partitioned into 0.1×0.1×0.1-μm^3^ transparent cubes. Synapse clusters were arrayed in a 3D hexagonal grid that was centered within the simulation volume and expanded or shrunk to vary spacing between synapses (Figure 7A,C; see Methods); to limit any synapse orientation bias, individual synapses were modeled without pre- and postsynaptic processes. Control simulations confirmed that this simplification did not affect iGluSnFR response amplitude or time course when imaged over ≥2 μm-radius ROIs (Supplementary Figure 2).

**Figure 7.**
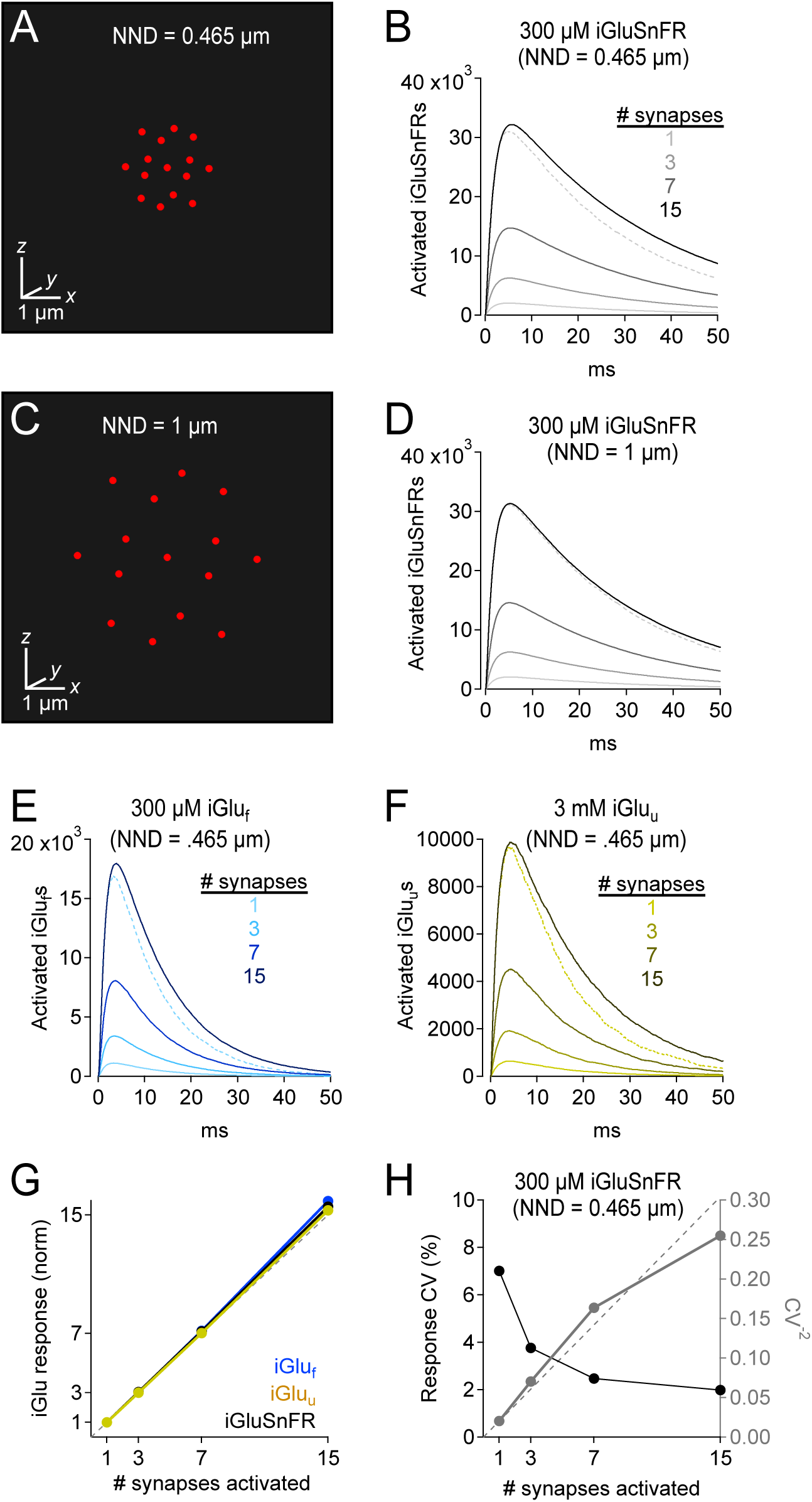
iGluSnFR and variants provide a linear indication of synaptic release. *A*, Schematic showing 15 active synapses clustered tightly (NND = 0.465 μm) in 3D diffusion space. *B*, Simulated iGluSnFR signals (300 μm iGluSnFR, 100 μm EAAT) elicited by coincident activation of 1, 3, 7, and 15 synapses (NND = 0.465 μm). Dotted gray trace shows the linear prediction for the 15-synapse case, i.e., the single synapse response multiplied by 15. *C* and *D*, As in *A* and *B*, but NND = 1 μm. *E*, As in *B* but for simulated iGlu_f_-mediated signals. *F*, As in *B* but for simulated iGlu_u_-mediated signals. *G*, Summary graph showing that iGlu signals provide a linear indication of synaptic release, even when NND = 0.465. H, iGluSnFR response variability (CV) also provides a reliable indication of relative numbers of activated synapses (see Faber and Korn, 1991).

Excitatory synapses are densely expressed in the CA1 region of the hippocampus, with a 465-nm average distance between nearest neighboring synapses (nearest neighbor distance, NND; Rusakov and Kullmann, 1998). When 15 simulated synapses were activated concomitantly at this density, iGluSnFR responses exhibited near perfect linearity, i.e., the compound response from 15 synapses was 15.6 times as large as the response from a single synapse (Figure 7B). Similar results were observed with iGlu_f_ and iGlu_u_ (Figure 7E-G). The slight supralinearity likely was due to slightly sublinear uptake reflecting a high degree of EAAT occupancy between activated synapses (data not shown).

Due to their extremely dense expression in astroglial membranes (Lehre and Danbolt, 1998), EAATs typically are not saturated even during trains of synaptic stimulation (Diamond and Jahr, 2000), enabling STCs to provide a linear indication of glutamate release (Diamond et al., 1998; Luscher et al., 1998). This is likely due to the fact that, because the release probability of individual CA1 synapses is ∼0.3 (Stevens and Wang, 1995; Hjelmstad et al., 1997), synaptic stimulation is unlikely to evoke coincident release at every synapse. Accordingly, when the NND of activated synapses was increased to 1 μm, simulated uptake rates and iGluSnFR responses exhibited near perfect linearity (Figure 7D; uptake data not shown).

Response variability is another way of measuring relative differences in the number of activated synapses (Faber and Korn, 1991). Specifically, if individual synapses exhibit similar binomial behavior to each other, the CV of the compound response will vary inversely with the square root of the number of activated synapses. Simulated iGluSnFR responses generally obeyed this relationship, as indicated in a plot of CV^-2^ vs. the number of activated synapses (Figure 7H). The slight deviation from the proportional relationship likely reflects subtle differences in quantal variability depending on the physical location of the synapse within the hexagonal array.

### Segregating EAATs and iGluSnFRs reduces buffering effects but does not speed iGluSnFR signal

It is possible that the buffering effects of iGluSnFR could be ameliorated by segregating EAATs and iGluSnFR into separate compartments, thereby giving EAATs some opportunity to take up glutamate without competing with iGluSnFR. This could be accomplished by, for example, expressing iGluSnFR only in some subset of interneuron plasma membranes. To test this idea, we confined EAATs and iGluSnFRs in simulations to alternating spherical shells of varying thickness, with EAATs occupying the innermost shell in each case (Figure 8A,B). EAAT and iGluSnFR concentrations within each shell were adjusted so that the average concentrations were 100 and 300 μM, respectively (Figure 8B). As expected, excluding iGluSnFR from the region immediately surrounding the release site reduced the amplitude of the indicator signal and sped the time course of glutamate uptake compared to the case in which iGluSnFRs and EAATs were perfectly co-localized (Figure 8C,D). Segregation did not, however, speed the iGluSnFR signal, because the subset of glutamate that diffused into the iGluSnFR-only region tended to remain there, buffered by the surrounding indicator and contributing to a prolonged iGluSnFR signal. This effect was also evident in the glutamate clearance time course, which exhibited a second, similarly slow component. The fast component of clearance, meanwhile, was similar to that observed in the absence of iGluSnFR (Figure 8D, dashed line). These results suggest that, when EAATs and iGluSnFRs are strongly segregated (e.g., 2-μm alternating shells), slow iGluSnFR signals may be observed even under conditions when most synaptically released glutamate is taken up quickly (red traces in Figure 8C,D). Similar results were obtained in simulations employing rectilinear coordinates with iGluSnFR and EAAT expression segregated alternately or randomly into 2×2×2 μm^3^ cubes (data not shown). These scenarios do not appear to reflect the experimental conditions reported above, however: iGluSnFR expression in astrocytic membranes slowed STCs significantly (Figure 3), whereas sharp segregation (2-μm shells or cubes) simulations produced STC waveforms similar in time course to those simulated in the absence of iGluSnFR (Figure 8E).

**Figure 8.**
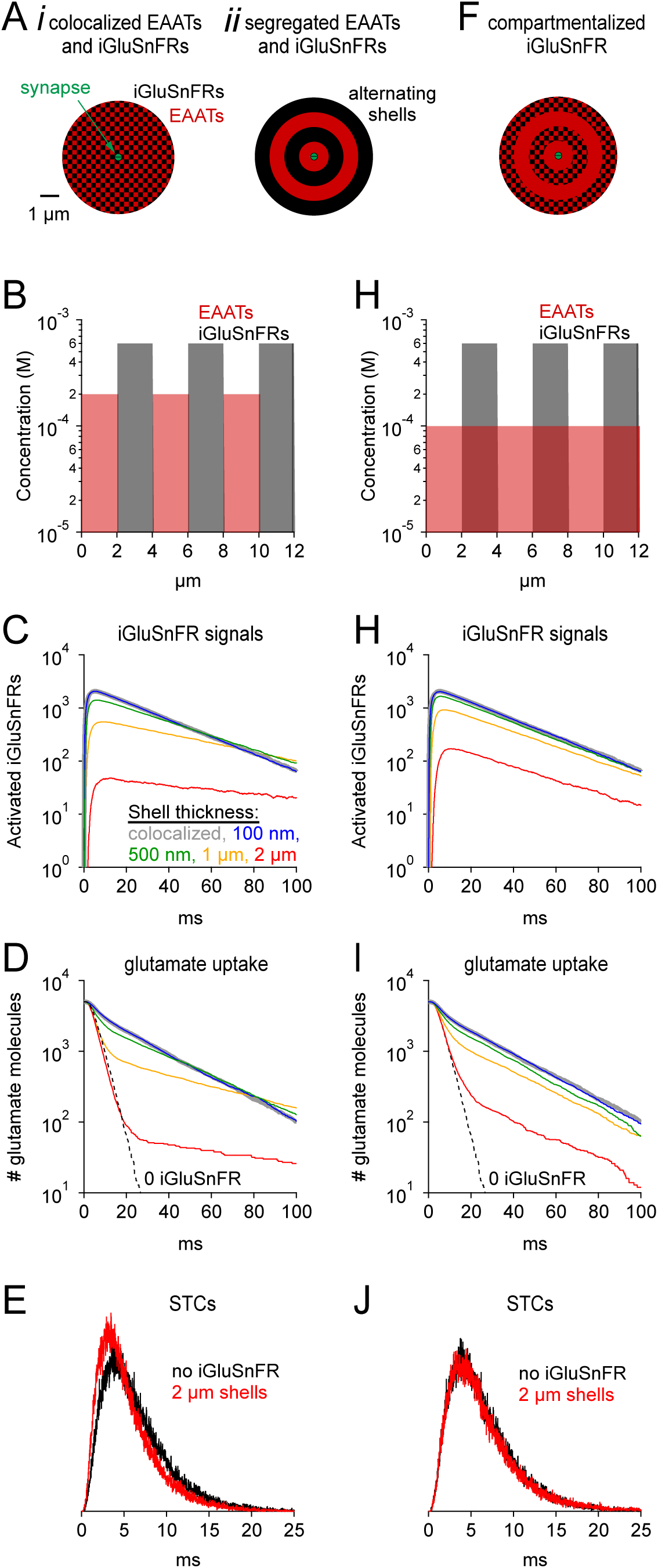
Segregated expression reduces buffering effects. *A*, Schematics of simulation in which EAATs and iGluSnFRs were colocalized (*i*), or segregated into alternating spherical shells surrounding the synapse, with EAATs occupying the innermost shell (*ii*). *B*, Concentration profile of EAATs and iGluSnFRs in the 2-μm shell case. In each case the average EAAT and iGluSnFR concentrations were 100 μM and 300 μM, respectively. *C*, Simulated iGluSnFR signal wave forms at four different shell thicknesses. *D*, Glutamate uptake time courses in the same simulations described in *C. E*, Simulated synaptic transporter currents (STCs) in the 2-μm shell condition and in the absence of iGluSnFR. *F-J*, As in *Aii-E*, except that EAATs were distributed evenly throughout all shells.

Finally, we tested a different scenario in which EAATs were expressed evenly in all shells and iGluSnFRs were expressed only in alternating shells (Figure 8F,G). This scenario may present a case in which astroglial EAATs sample extracellular space evenly but iGluSnFR is expressed in only a subset of neurons. In this case, iGluSnFR signals were larger and faster than in the segregated case, and clearance was also faster (cf. Figure 8C,D and H,I). The iGluSnFR signals were larger because the same EAAT concentration, evenly distributed, presented half the EAATs in the initial shell, enabling more glutamate to reach the iGluSnFR-containing shells. iGluSnFR signals and glutamate clearance were faster because glutamate no longer became “trapped” in iGluSnFR-only regions. Nonetheless, the simulated STC was very similar to that in the absence of iGluSnFR (Figure 8J). Similar results were obtained using Cartesian coordinates (data not shown). These results indicate that iGluSnFR, when expressed in only a subset of neuronal membranes, may produce detectable (albeit smaller) signals with reduced distortion of endogenous glutamate uptake dynamics. Such an arrangement may be ideal for experiments in which many synapses can be imaged simultaneously but glutamate diffusion and uptake must not be disturbed.

## Discussion

iGluSnFR and its variants offer great opportunities to study the dynamics of excitatory synaptic transmission in the brain. The diffusion simulations presented here aimed to examine what factors influence the measured time course of iGluSnFR signals. They suggest that iGluSnFR responses are slower than predicted from diffusion laws and astrocyte STC recordings because iGluSnFR buffers glutamate diffusion and prolongs its extracellular lifetime following synaptic release. Consistent with these conclusions, STCs recorded from iGluSnFR^+^ cortical astrocytes were slower than those recorded in tdTomato^+^ control astrocytes, confirming that iGluSnFR slows glutamate clearance. The buffering effects of iGluSnFR were also evident in simulations incorporating lower affinity iGluSnFR variants. Compartmentalized iGluSnFR expression, perhaps in a subset of neuronal membranes, might reduce the buffering effect but likely would not produce faster signals that more closely approximated the native glutamate clearance time course. Despite these caveats regarding time course, multi-synapse simulations suggest that glutamate indicators are linear indicators of the number of activated synapses.

### Indicator expression levels influence glutamate uptake and iGluSnFR signal time courses

Our simulations clearly indicate that the buffering effects and slowed glutamate clearance depend strongly on the expression levels of iGluSnFR (Figure 2), potentially complicating comparison of iGluSnFR signals across brain regions. For example, a recent report indicate that iGluSnFR signals are faster in the hippocampus than the cortex, suggesting that glutamate uptake in the hippocampus is more efficient (Pinky et al., 2018). Absent other considerations, these differences could reflect differences in iGluSnFR expression rather than uptake capacity, i.e., it may be that iGluSnFR is simply expressed more densely in the cortex, thereby slowing iGluSnFR signals recorded there (Figure 2C). In this particular case, however, STCs are also faster in hippocampus than cortex (Hanson et al., 2015), providing a second, complementary test that corroborates the first. Importantly, these STC recordings were made from astrocytes in the *absence* of iGluSnFR expression (Hanson et al., 2015): Our simulations and experiments indicate that STCs recorded in iGluSnFR^+^ tissue also are slowed relative to control, potentially providing a misleading agreement between STCs and iGluSnFR fluorescence signals. These results also suggest that iGlu variants optimized for increased expression (Marvin et al., 2018), actually may disrupt glutamatergic signaling even more.

iGluSnFR concentration is likely the most critical parameter in our simulations that is not constrained in some way by experimental data. The actual effective concentration likely varies widely due to differences in brain region, expression system, promotor and indicator subtype. Biochemical measures may overestimate this parameter by including protein that is not expressed in the plasma membrane, and immunohistochemistry cannot distinguish what fraction of molecules on the cell surface is active. In the case of EAATs, quantitative immunoblotting revealed extremely high endogenous expression of EAAT1 and EAAT2 in the hippocampus and cerebellum (Lehre and Danbolt, 1998), and postembedding immunoelectron microscopy indicated that most EAATs are localized to astrocytic plasma membranes (Chaudhry et al., 1995). EAAT expression is particularly dense in the hippocampus (Lehre et al., 1995), exceeding 10,000 monomers per μm^2^ of astroglial plasma membrane (Lehre and Danbolt, 1998). It is, admittedly, remarkable that iGluSnFR might be expressed at even higher levels, as predicted by our simulations. It also is unknown whether iGluSnFR expression in astrocytes might come at the expense of surface EAAT expression, although cortical astrocyte STCs recorded from tissue in which iGluSnFR is expressed in neurons are similar in waveform to those recorded here in iGluSnFR-expressing astrocytes (Figure 3; M. Armbruster and C.G. Dulla, unpublished observations).

### Effects on extrasynaptic signaling

Our simulations predicted that iGluSnFR reduces the free concentration of extrasynaptic glutamate (Figure 2G,H) and may therefore influence the actions of synaptically released glutamate on extrasynaptic metabotropic receptors, or perhaps glutamate spillover between excitatory synapses (Arnth-Jensen et al., 2002; Scimemi et al., 2004), thereby influencing aspects of synaptic signaling and plasticity.

Extrasynaptic buffering may play a particularly significant role if the target neurotransmitter typically acts at receptors located some distance from its release site. For example, dense expression of indicators for dopamine (Sun et al., 2018), norepinephrine, or serotonin might influence substantially the modulatory effects of those transmitters, although dopamine has been shown to act, at least in some cases, more locally than previously expected (Courtney and Ford, 2014).

### Model Limitations

The Monte Carlo diffusion model used here includes significant simplifications that dramatically reduce the computational resources required but may also compromise somewhat the accuracy of the results. The choice to model the extrasynaptic space as an isotropic region (Rusakov and Kullmann, 1998; Diamond, 2005), rather than instantiating an explicit structure (Mishchenko et al., 2010), allows the model of a single synapse to represent the average arrangement across many synapses. EAATs and iGluSnFRs were evenly distributed within spherical or cubic partitions surrounding the synapse, regular arrangements that surely differ from the endogenous structure. Similar results were obtained when partition thickness was varied from zero (i.e., continuous) to 500 nm (spherical example shown in Figure 8), suggesting that abstracting the fine structure of the extracellular space did not influence the results significantly. The spherical ROIs used here likely overestimates background fluorescence (F_0_) in experiments using line scans across individual synapses (Helassa et al., 2018). ΔF/F_0_ values were considered here primarily to compare different simulation parameters, as experimentally observed ΔF/F_0_ values (and indicator affinity) are likely to vary with experimental and imaging conditions (Marvin et al., 2013; Helassa et al., 2018; Marvin et al., 2018).

Explicitly modeling only those EAATs and iGluSnFRs that bind synaptically released glutamate drastically reduced the computational power required to simulate very high levels of EAAT/iGluSnFR expression over large volumes (Diamond, 2005). For example, explicitly simulating 3 mM iGluSnFR in a 30×30×30 μm^3^ volume would require tracking almost 1.5×10^11^ Markov states (i.e., ∼150 GB of RAM for iGluSnFR alone), as opposed to a maximum here of 2.2×10^5^ (simulating 15 synapses releasing a total of 75,000 glutamate molecules; Figure 7). It did reduce binding interactions to simple probabilities (Methods) at the expense of greater detail in more computationally extensive simulations (Stiles and Bartol, 2001) and would, therefore be insufficient to simulate steric ligand-receptor interactions or competition between individual, adjacent receptors.

### Is there an ideal glutamate indicator?

Efforts to optimize glutamate indicators generally aim to make them faster, brighter, or express more strongly (Helassa et al., 2018; Marvin et al., 2018). Our simulations suggest that increasing expression could further disrupt glutamate diffusion (Figure 2C) and, due to increased background fluorescence, may actually decrease ΔF/F_0_ (Figure 4). To examine what kinetic properties would yield the best performance, we combined the most advantageous kinetic properties of iGlus into one hypothetical indicator (FrankenSnFR; Supplementary Figure 3A). A fast unbinding rate (k_-1_) ensures rapid deactivation but necessitates a fast binding rate (k_+1_) to maintain suitably high affinity and rapid responses at low expression levels. Both the entry and exit from the activated state (k_+2_ and k_-2_, respectively) must be fast to preserve rapid signal onset and cessation, and k_+2_ must be significantly greater than k_-2_ to achieve a high maximal activation probability (P_max_ = k_+2_/[k_+2_ + k_-2_]). Combining these features created an indicator that activated and deactivated rapidly, bound glutamate with high affinity (k_d_ = 71 μM, EC_50_=15 μM) and exhibited high P_max_ (0.77; Supplementary Figure 3B-D). FrankenSnFR exhibited reasonable single synapse activation at low expression levels (e.g., 3 μM, Supplementary Figure 3E) without disrupting glutamate clearance (Supplementary Figure 3F). Similar results were observed with ≤10 μM iGluSnFR (Figure 2), but FrankenSnFR delivered much faster response kinetics (Supplementary Figure 3E). Nonetheless, at higher expression levels FrankenSnFR disrupted diffusion and uptake just like the others. These simulations suggest that the greatest improvements in iGlu performance are gained through increasing P_max_ and, of course, dynamic range (ΔF/F_0_), so that high signal-to-noise characteristics can be achieved at low expression levels.

## Methods

All animal protocols were approved by the Tufts Institutional Animal Care and Use Committee.

### Adeno-associated virus injection

C57BL/6 male and female mice (P30-35) were stereotaxtically injected with either GFAP-iGluSnFR or GfaABC1D-tdtomato (University of Pennsylvania Vector Core; catalog #AV-5-PV2723, AV-5-PV3106) in a single hemisphere with 3 injections sites (coordinates): (1.25, 1.25, 0.5), (1.25, 2.25, 0.5), and (1.25, 3.25, 0.5) (λ + *x*, +*y*, −*z*) mm. Mice were anesthetized with isoflurane for surgery, reporter viruses were injected (1 μL per site, 0.15 μL/min) with ∼5×10^9^ gene copies. Mice were housed in 12/12 light/dark cycles following surgeries and were used for acute slice preparations 21–28 d following injection.

### Preparation of acute brain slices

Cortical brain slices were prepared from tdTomato or iGluSnFR-infected C57/B6 mice (Armbruster et al., 2016). Mice were anesthetized with isoflurane, decapitated, and the brains were rapidly removed and placed in ice-cold slicing solution containing (in mM): 2.5 KCl, 1.25 NaH_2_PO_4_, 10 MgSO_4_, 0.5 CaCl_2_, 11 glucose, 234 sucrose, and 26 NaHCO_3_ and equilibrated with 95% O_2_:5% CO_2_. The brain was glued to a Vibratome VT1200S (Leica Microsystems, Wetzlar, Germany), and slices (400 μm thick) were cut in a coronal orientation. Slices were then placed into a recovery chamber containing aCSF comprising (in mM): 126 NaCl, 2.5 KCl, 1.25 NaH_2_PO_4_, 1 MgSO_4_, 2 CaCl_2_, 10 glucose, and 26 NaHCO_3_ (equilibrated with 95% O_2_:5% CO_2_). Slices were allowed equilibrate in aCSF at 32°C for 1 h. Slices were loaded with sulforhodamine 101 (SR-101, 0.5 μM) in aCSF for 5 min at 32°C before equilibration (Nimmerjahn et al., 2004) and were allowed to return to room temperature prior to electrophysiology/imaging.

### Glutamate transporter currents

Glutamate transporter currents were recorded similarly to previous studies (Diamond, 2005; Armbruster et al., 2016). Acute slices were placed into a submersion chamber (Warner Instruments, Hamden, CT), held in place with small gold wires, and perfused with aCSF containing DNQX (20 μM), AP5 (50 μM) and Gabazine (SR95531, 10 μM) to block AMPA, NMDA and GABA_A_ receptors, respectively. Additionally, aCSF contained BaCl_2_ (200 μM) to block astrocyte K^+^ conductances and isolate transporter currents (Ransom and Sontheimer, 1995; Afzalov et al., 2013; Armbruster et al., 2016). aCSF was equilibrated with 95% O_2_:5% CO_2_ and circulated at 2 ml/min at 34°C. A tungsten concentric bipolar stimulating electrode (FHC) was placed in the deep cortical layers, and astrocytes were patched and/or iGluSnFR imaged in layer II/III. Astrocytes were identified by morphology (small, round cell bodies), membrane properties, and SR-101/tdTomato labeling as imaged with a Cy3 filter cube (excitation 560/40 nm, emission 630/75 nm, Chroma). Astrocyte internal solution contained the following (in mM): 120 potassium gluconate, 20 HEPES, 10 EGTA, 2 MgATP, and 0.2 NaGTP. 4–12 MΩ borosilicate pipettes were used to establish whole-cell patch-clamp recordings using a Multiclamp 700B patch-clamp amplifier, sampled at 10 kHz using pClamp software (Molecular Devices, San Jose, CA). Once a whole-cell recording was established, cells were confirmed as astrocytes based on their passive membrane properties, low membrane resistance, and hyperpolarized resting membrane potential.

100 μs stimulus pulses were generated every 15 s with a stimulus isolator (ISO-Flex, A.M.P.I., Jerusalem, Israel). Stimulus intensity was set at 2× the resolvable threshold stimulation. Simultaneous electrophysiology and iGluSnFR imaging were performed in a subset of cells. Imaging was performed using a Prime95b (Teledyne Photometrics, Tucson, AZ) camera with a 200Hz frame rate, illuminated by a CoolLED illuminator and GFP filter cube (Chroma Technology, Bellows Falls, VT) and controlled by MicroManager (Edelstein et al., 2014) with a 60× water-immersion objective (LUMPLANFL, Olympus, Waltham, MA) on an Olympus/Prior Openscope microscope. The imaged region was 97 × 37 μm^2^ (530 × 200 pixels, 183 nm per pixel).

### Analysis

Analysis was performed using MATLAB (The MathWorks, Natick, MA) and Origin (Originlab, Northampton, MA). For astrocyte synaptic transporter current recordings, 4–12 sweeps were averaged and normalized, and the decay of the glutamate transporter current was fit with a mono-exponential function (plus *y* offset) to quantify glutamate uptake kinetics (fitting region was 18-148 ms post-stimulus). For iGluSnFR imaging, 10 repeated runs of identical stimulation were averaged together and decays were fit with a bi-exponential function (decay + bleaching).

### Simulations

Transmitter diffusion, uptake and glutamate indicator activation were simulated using an expanded version of a previous model (Diamond, 2005) written in MATLAB. Results were analyzed and graphed using IgorPro (WaveMetrics, Lake Oswego, OR).

Glutamate diffusion in single-synapse simulations was modeled as a random walk of 5000 independent glutamate molecules originating simultaneously from a point source in the center of a 320-nm-diameter, 20-nm thick synaptic cleft (Ventura and Harris, 1999). At each time step Δt = 1 µs (Δt = 0.5 or 2 µs yielded similar results), each glutamate molecule was displaced in each spatial dimension by a distance r randomly selected from a normal distribution about zero (average r^2^=2DΔt; Hille, 1984, where D (the diffusion coefficient) = 0.253 µs^2^ ms^-1^ in extracellular fluid at 25°C; Longsworth, 1953; Nielsen et al., 2004). Diffusion within the synaptic cleft was limited to two (*x,y*) dimensions (Barbour and Hausser, 1997); extrasynaptic diffusion was modeled three-dimensionally through an isotropic extrasynaptic space (extracellular volume fraction = 0.21). D in all spaces was reduced further to account for tortuosity of the extracellular space (D^*^ = D/λ^2^; λ = 1.55; Rusakov and Kullmann, 1998; eliminating tortuosity within the synaptic cleft did not affect the results significantly). For multi-synapse simulations (Figure 6), synapses were modeled as point sources within isotropic extracellular space. Control simulations confirmed that removal of the synaptic cleft had little effect on the simulated iGluSnFR waveform for ROIs > 2 μm radius (Supplemental Figure 2).

Transporters were modeled using a Markov representation of EAAT2 (Bergles et al., 2002; Supplemental Figure 1A), with two simplifying modifications: The extracellular transporter was configured to bind H^+^ prior before glutamate, rather than allowing either to bind first, and the T_i_Na_2_⇋T_o_Na_2_ transition was eliminated. Once all of the transported elements unbound on the intracellular side, the glutamate molecule was designated as taken up and removed from the simulation and the transporter returned to the unbound, outward facing state at a rate corresponding to physiological measured recovery rate (Bergles et al., 2002). Substrate concentrations other than [glu]_o_ were assumed constant. Simulated STC waveforms (Figure 2B, inset, Figure 8E,J) reflected the stoichiometric current in the model (+1 for forward transitions 1, 7 and 9, −1 for forward transition 15 in Bergles et al., 2002). Voltage-dependent rates were calculated at −95 mV (similar results were observed at −70 mV, data not shown). iGluSnFR kinetics were implemented according to a simple three-state model: bound, unbound, and fluorescent (Helassa et al., 2018; Supplementary Figure 1B).

Extracellular space was partitioned transparently into 10-nm-think concentric spherical shells (single-synapse simulations) or 100×100×100 nm^3^ cubes (multi-synapse simulations) so that local transporter, iGluSnFR and glutamate concentration could be determined. At each time step, the probability of binding to a transporter or iGluSnFR was determined independently for each glutamate molecule as follows: First, the EAAT2 and iGluSnFR glutamate binding rates (Bergles et al., 2002; Helassa et al., 2018) were multiplied by the time step, the glutamate concentration in the relevant shell/cube and the number of transporter/iGluSnFR molecules in the shell/cube, to give the number of transporters/iGluSnFRs bound in the time step, and then divided by the number of glutamate molecules in the shell/cube to yield the probability that a particular glutamate molecule would bind. If binding occurred (i.e., if a random number between 0 and 1 was less than the binding probability), the number of free transporter/iGluSnFR molecules in the cell was decremented. Once bound, the glutamate molecule underwent probabilistic transitions in subsequent time steps through the Markov schemes. Because transporters and iGluSnFRs were modeled explicitly only upon binding an individual glutamate molecule, the number of simulated transporters/iGluSnFRs was limited by the relatively low number (5000-75,000) of glutamate molecules simulated.

## Acknowledgements

This work was supported by the NIH (NS113499, NS104478, NS100796 to C.G.D.) and the NINDS Intramural Research Program (NS003039 to J.S.D.). We thank David DiGregorio for helpful comments on the manuscript. The authors have no competing interests to disclose.

## FIGURE LEGENDS

**Supplementary Figure 1 | Kinetic models.** *A*, Multi-state model of EAAT2 used in the simulations (Bergles et al., 2002). *B*, Multi-state models used to simulate iGluSnFR, iGlu_f_ and iGlu_u_ (Helassa et al., 2018).

**Supplementary Figure 2 | Including the synaptic cleft has little effect on simulated iGluSnFR time courses.** *A*, Simulated iGluSnFR signals in a radially symmetric model that either included (blue) or excluded (gold) a 320-nm-diameter synaptic cleft. *B*, Summary of amplitudes of simulated iGluSnFR signals with and without an explicitly modeled synaptic cleft. *C*, As in B but showing exponential decay times. *D*, Comparison of glutamate uptake time course.

**Supplementary Figure 3 | Characteristics of a theoretically ideal glutamate indicator.** *A*, Rate constants (see state diagram in Supplementary Figure 1B) used to create FrankenSnFR, a hypothetical glutamate indicator. B, Simulated FrankenSnFR activation by 20-ms applications of glutamate (concentrations varied logarithmically from 1 μM to 30 mM). *C*, Comparison of simulated glutamate dose-response curves for iGluSnFR (black), iGlu_f_ (blue), iGlu_u_ (gold) and FrankenSnFR (green). *D*, Responses of indicators (same color scheme as *C*) to 1 mM glutamate, normalized and superimposed to compare activation and deactivation kinetics. *E*, Simulated FrankenSnFR responses (three different concentrations, spherical ROI radius = 10 μm) to the synaptic release of 5000 glutamate molecules. Response of 10 μM iGluSnFR shown for comparison (dashed black trace). *F*, Glutamate uptake time course in the simulations shown in *E*, as well as clearance in the absence of any indicator (gray).

## References

Afzalov R, Pryazhnikov E, Shih PY, Kondratskaya E, Zobova S, Leino S, Salminen O, Khiroug L, Semyanov A (2013) Low micromolar Ba(2+) potentiates glutamate transporter current in hippocampal astrocytes. Front Cell Neurosci 7:135.

Armbruster M, Hanson E, Dulla CG (2016) Glutamate Clearance Is Locally Modulated by Presynaptic Neuronal Activity in the Cerebral Cortex. Journal of Neuroscience 36:10404–10415.

Arnth-Jensen N, Jabaudon D, Scanziani M (2002) Cooperation between independent hippocampal synapses is controlled by glutamate uptake. Nature Neuroscience 5:325–331.

Barbour B (2001) An evaluation of synapse independence. Journal of Neuroscience 21:7969–7984.

Barbour B, Hausser M (1997) Intersynaptic diffusion of neurotransmitter. Trends Neurosci 20:377–384.

Bergles DE, Jahr CE (1997) Synaptic activation of glutamate transporters in hippocampal astrocytes. Neuron 19:1297–1308.

Bergles DE, Tzingounis AV, Jahr CE (2002) Comparison of coupled and uncoupled currents during glutamate uptake by GLT-1 transporters. Journal of Neuroscience 22:10153–10162.

Borghuis BG, Marvin JS, Looger LL, Demb JB (2013) Two-photon imaging of nonlinear glutamate release dynamics at bipolar cell synapses in the mouse retina. Journal of Neuroscience 33:10972–10985.

Chaudhry FA, Lehre KP, van Lookeren Campagne M, Ottersen OP, Danbolt NC, Storm-Mathisen J (1995) Glutamate transporters in glial plasma membranes: highly differentiated localizations revealed by quantitative ultrastructural immunocytochemistry. Neuron 15:711–720.

Courtney NA, Ford CP (2014) The timing of dopamine- and noradrenaline-mediated transmission reflects underlying differences in the extent of spillover and pooling. Journal of Neuroscience 34:7645–7656.

de Lorimier R, Smith J, Dwyer M, Looger L, Sali K, Paavola C, Rizk S, Sadigov S, Conrad D, Loew L, Helinga H (2002) Construction of a fluorescent biosensor family. Protein Science 11:2655–2675.

Diamond JS (2005) Deriving the glutamate clearance time course from transporter currents in CA1 hippocampal astrocytes: transmitter uptake gets faster during development. Journal of Neuroscience 25:2906–2916.

Diamond JS, Jahr CE (1997) Transporters buffer synaptically released glutamate on a submillisecond time scale. Journal of Neuroscience 17:4672–4687.

Diamond JS, Jahr CE (2000) Synaptically released glutamate does not overwhelm transporters on hippocampal astrocytes during high-frequency stimulation. J Neurophysiol 83:2835–2843.

Diamond JS, Bergles DE, Jahr CE (1998) Glutamate release monitored with astrocyte transporter currents during LTP. Neuron 21:425–433.

Edelstein AD, Tsuchida MA, Amodaj N, Pinkard H, Vale RD, Stuurman N (2014) Advanced methods of microscope control using muManager software. J Biol Methods 1.

Faber DS, Korn H (1991) Applicability of the coefficient of variation method for analyzing synaptic plasticity. Biophysical Journal 60:1288–1294.

Fehr M, Frommer WB, Lalonde S (2002) Visualization of maltose uptake in living yeast cells by fluorescent nanosensors. Proc Natl Acad Sci U S A 99:9846–9851.

Franke K, Berens P, Schubert T, Bethge M, Euler T, Baden T (2017) Inhibition decorrelates visual feature representations in the inner retina. Nature 542:439–444.

Hanson E, Armbruster M, Cantu D, Andresen L, Taylor A, Danbolt NC, Dulla CG (2015) Astrocytic glutamate uptake is slow and does not limit neuronal NMDA receptor activation in the neonatal neocortex. Glia 63:1784–1796.

Helassa N, Durst CD, Coates C, Kerruth S, Arif U, Schulze C, Wiegert JS, Geeves M, Oertner TG, Torok K (2018) Ultrafast glutamate sensors resolve high-frequency release at Schaffer collateral synapses. Proc Natl Acad Sci U S A 115:5594–5599.

Hille B (1984) Ionic Channels of Excitable Membranes, 1st Edition. Sunderland, Massachusetts: Sinauer Associates, Inc.

Hires SA, Zhu Y, Tsien RY (2008) Optical measurement of synaptic glutamate spillover and reuptake by linker optimized glutamate-sensitive fluorescent reporters. Proc Natl Acad Sci U S A 105:4411–4416.

Hjelmstad GO, Nicoll RA, Malenka RC (1997) Synaptic refractory period provides a measure of probability of release in the hippocampus. Neuron 19:1309–1318.

Lehre KP, Danbolt NC (1998) The number of glutamate transporter subtype molecules at glutamatergic synapses: chemical and stereological quantification in young adult rat brain. Journal of Neuroscience 18:8751–8757.

Lehre KP, Levy LM, Ottersen OP, Storm-Mathisen J, Danbolt NC (1995) Differential expression of two glial glutamate transporters in the rat brain: quantitative and immunocytochemical observations. Journal of Neuroscience 15:1835–1853.

Longsworth L (1953) Diffusion measurements at 25º of aqueous solutions of amino acids, peptides and sugars. J Am Chem Soc 75:5705–5709.

Luscher C, Malenka RC, Nicoll RA (1998) Monitoring glutamate release during LTP with glial transporter currents. Neuron 21:435–441.

Marvin JS, Borghuis BG, Tian L, Cichon J, Harnett MT, Akerboom J, Gordus A, Renninger SL, Chen TW, Bargmann CI, Orger MB, Schreiter ER, Demb JB, Gan WB, Hires SA, Looger LL (2013) An optimized fluorescent probe for visualizing glutamate neurotransmission. Nat Methods 10:162–170.

Marvin JS et al. (2018) Stability, affinity, and chromatic variants of the glutamate sensor iGluSnFR. Nat Methods 15:936–939.

Mishchenko Y, Hu T, Spacek J, Mendenhall J, Harris KM, Chklovskii DB (2010) Ultrastructural analysis of hippocampal neuropil from the connectomics perspective. Neuron 67:1009–1020.

Nicholson C, Sykova E (1998) Extracellular space structure revealed by diffusion analysis. Trends Neurosci 21:207–215.

Nielsen TA, DiGregorio DA, Silver RA (2004) Modulation of glutamate mobility reveals the mechanism underlying slow-rising AMPAR EPSCs and the diffusion coefficient in the synaptic cleft. Neuron 42:757–771.

Nimmerjahn A, Kirchhoff F, Kerr JN, Helmchen F (2004) Sulforhodamine 101 as a specific marker of astroglia in the neocortex in vivo. Nat Methods 1:31–37.

Okumoto S, Looger LL, Micheva KD, Reimer RJ, Smith SJ, Frommer WB (2005) Detection of glutamate release from neurons by genetically encoded surface-displayed FRET nanosensors. Proc Natl Acad Sci U S A 102:8740–8745.

Parsons MP, Vanni MP, Woodard CL, Kang R, Murphy TH, Raymond LA (2016) Real-time imaging of glutamate clearance reveals normal striatal uptake in Huntington disease mouse models. Nature Communications 7:11251.

Pinky NF, Wilkie CM, Barnes JR, Parsons MP (2018) Region- and Activity-Dependent Regulation of Extracellular Glutamate. Journal of Neuroscience 38:5351–5366.

Quiocho FA, Spurlino JC, Rodseth LE (1997) Extensive features of tight oligosaccharide binding revealed in high-resolution structures of the maltodextrin transport/chemosensory receptor. Structure 5:997–1015.

Ransom CB, Sontheimer H (1995) Biophysical and pharmacological characterization of inwardly rectifying K+ currents in rat spinal cord astrocytes. J Neurophysiol 73:333–346.

Rusakov DA, Kullmann DM (1998) Extrasynaptic glutamate diffusion in the hippocampus: ultrastructural constraints, uptake, and receptor activation. Journal of Neuroscience 18:3158–3170.

Schikorski T, Stevens CF (1997) Quantitative ultrastructural analysis of hippocampal excitatory synapses. Journal of Neuroscience 17:5858–5867.

Scimemi A, Fine A, Kullmann DM, Rusakov DA (2004) NR2B-containing receptors mediate cross talk among hippocampal synapses. Journal of Neuroscience 24:4767–4777.

Stevens CF, Wang Y (1995) Facilitation and depression at single central synapses. Neuron 14:795–802.

Stiles J, Bartol T (2001) Monte Carlo methods for simulating realistic synaptic microphysiology using MCell. In: Computational Neuroscience: Realistic Modeling for Experimentalists (De Schutter E, ed), pp 87–127. Boca Raton: CRC Press.

Sun F et al. (2018) A Genetically Encoded Fluorescent Sensor Enables Rapid and Specific Detection of Dopamine in Flies, Fish, and Mice. Cell 174:481–496 e419.

Thomas CG, Tian H, Diamond JS (2011) The relative roles of diffusion and uptake in clearing synaptically released glutamate change during early postnatal development. Journal of Neuroscience 31:4743–4754.

Ventura R, Harris KM (1999) Three-dimensional relationships between hippocampal synapses and astrocytes. Journal of Neuroscience 19:6897–6906.

Wadiche JI, Amara SG, Kavanaugh MP (1995) Ion fluxes associated with excitatory amino acid transport. Neuron 15:721–728.

Wahl LM, Pouzat C, Stratford KJ (1996) Monte Carlo simulation of fast excitatory synaptic transmission at a hippocampal synapse. J Neurophysiol 75:597–608.

Yonehara K, Farrow K, Ghanem A, Hillier D, Balint K, Teixeira M, Juttner J, Noda M, Neve RL, Conzelmann KK, Roska B (2013) The first stage of cardinal direction selectivity is localized to the dendrites of retinal ganglion cells. Neuron 79:1078–1085.

