## Supplementary Figures 1-3 for "Effects of fluorescent glutamate indicators on neurotransmitter diffusion and uptake"

### Supplementary Figure 1

#### A Glutamate transporter model

$$T_oK \overset{16}{\rightleftharpoons} T_o \overset{1}{\rightleftharpoons} T_oNa_1 \overset{2}{\rightleftharpoons} T_oNa_2 \overset{4}{\rightleftharpoons} T_oNa_2H \overset{6}{\rightleftharpoons} T_oNa_2GH \overset{7}{\rightleftharpoons} T_oNa_3GH$$

$$15 \upharpoonleft\!\downharpoonright \qquad\qquad\qquad\qquad\qquad\qquad\qquad\qquad \upharpoonleft\!\downharpoonright 8$$

$$T_iK \underset{14}{\rightleftharpoons} T_i \underset{13}{\rightleftharpoons} T_iNa_1 \underset{12}{\rightleftharpoons} T_iNa_2 \underset{11}{\rightleftharpoons} T_iNa_2G \underset{10}{\rightleftharpoons} T_iNa_2GH \underset{9}{\rightleftharpoons} T_iNa_3GH$$

| | Forward (↻) | Backward (↺) | | Forward (↻) | Backward (↺) |
|---|---|---|---|---|---|
| 1 | $1\times10^{4}$ $M^{-1}s^{-1}$ | 100 $s^{-1}$ | 10 | 3000 $s^{-1}$ | $9\times10^{10}$ $M^{-1}s^{-1}$ |
| 2 | $1\times10^{4}$ $M^{-1}s^{-1}$ | 500 $s^{-1}$ | 11 | 3000 $s^{-1}$ | $1\times10^{5}$ $M^{-1}s^{-1}$ |
| 4 | $6\times10^{11}$ $M^{-1}s^{-1}$ | 700 $s^{-1}$ | 12 | $1\times10^{5}$ $s^{-1}$ | $2\times10^{7}$ $M^{-1}s^{-1}$ |
| 6 | $6\times10^{6}$ $M^{-1}s^{-1}$ | 500 $s^{-1}$ | 13 | $1\times10^{5}$ $s^{-1}$ | $1\times10^{8}$ $M^{-1}s^{-1}$ |
| 7 | $1\times10^{4}$ $M^{-1}s^{-1}$ | 1000 $s^{-1}$ | 14 | $1\times10^{6}$ $M^{-1}s^{-1}$ | 1000 $s^{-1}$ |
| 8 | 2000 $s^{-1}$ | 1900 $s^{-1}$ | 15 | 40$s^{-1}$ | 10 $s^{-1}$ |
| 9 | 1000 $s^{-1}$ | $4\times10^{4}$ $M^{-1}s^{-1}$ | 16 | $2\times10^{4}$ $s^{-1}$ | $1\times10^{6}$ $M^{-1}s$ |

#### B iGluSnFR models

$$S \underset{k_{-1}}{\overset{k_{+1}}{\rightleftharpoons}} SG \underset{k_{-2}}{\overset{k_{+2}}{\rightleftharpoons}} SG^{*}$$

| | iGluSnFR | $iGlu_f$ | $iGlu_u$ |
|---|---|---|---|
| $k_{+1}$ | $2.7\times10^{7}$ $M^{-1}s^{-1}$ | $3.5\times10^{6}$ $M^{-1}s^{-1}$ | $2.2\times10^{6}$ $M^{-1}s^{-1}$ |
| $k_{-1}$ | 5965 $s^{-1}$ | 2206 $s^{-1}$ | 1704 $s^{-1}$ |
| $k_{+2}$ | 569 $s^{-1}$ | 944 $s^{-1}$ | 136 $s^{-1}$ |
| $k_{-2}$ | 110 $s^{-1}$ | 283 $s^{-1}$ | 468 $s^{-1}$ |
| $F_{on}$ | 25.4 | 19.4 | 15.3 |
| $F_{off}$ | 6.1 | 5.7 | 5.3 |

### Supplementary Figure 2

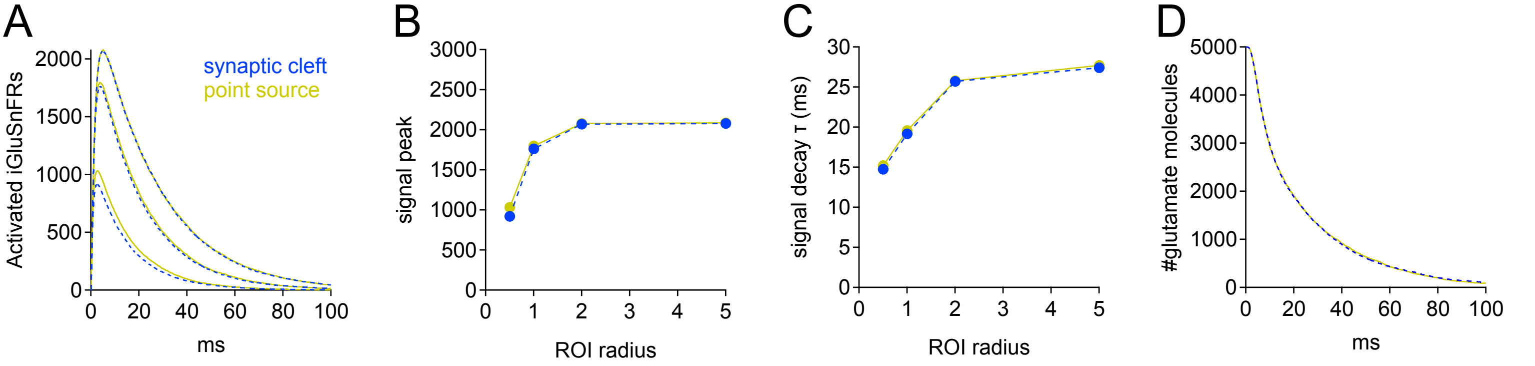

### Supplementary Figure 3

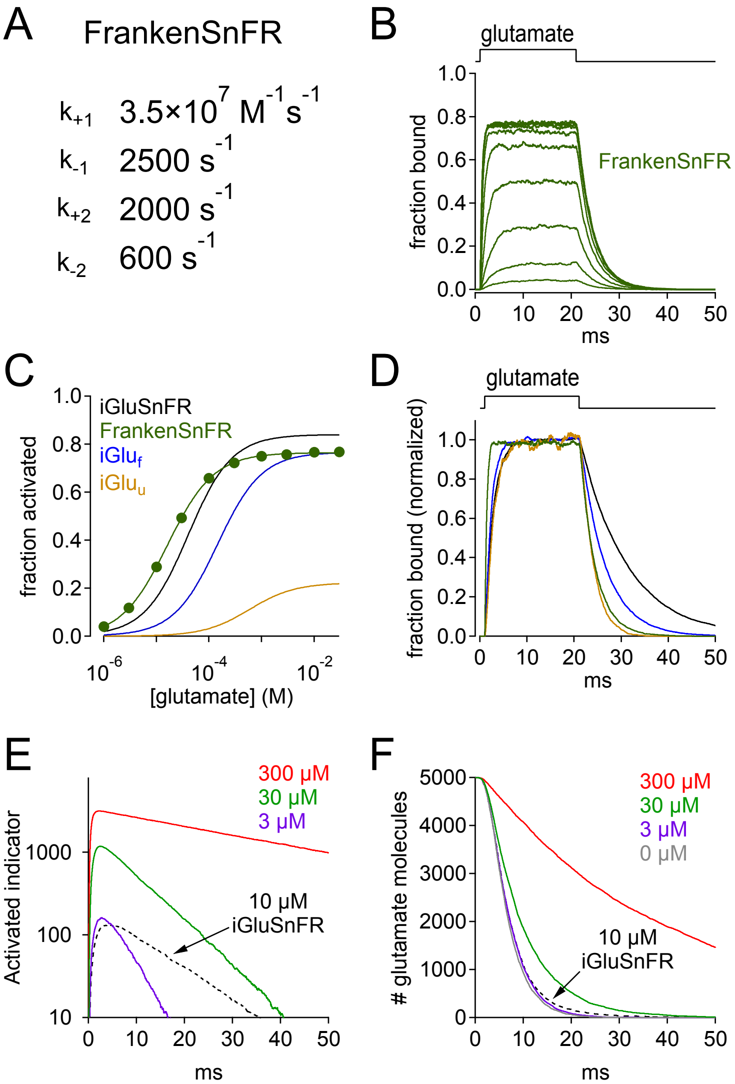
